# Neuromodulatory effects on synchrony and network reorganization in networks of coupled Kuramoto oscillators

**DOI:** 10.1101/2024.02.27.582261

**Authors:** Sinan Aktay, Leonard M. Sander, Michal Zochowski

**Affiliations:** Biophysics Program, University of Michigan, Ann Arbor, Michigan 48105, USA; Department of Physics, University of Michigan, Ann Arbor, Michigan 48105, USA

## Abstract

Neuromodulatory processes in the brain can critically change signal processing on a cellular level leading to dramatic changes in network level reorganization. Here, we use coupled non-identical Kuramoto oscillators to investigate how changes in the shape of phase response curves from Type 1 to Type 2, mediated by varying ACh levels, coupled with activity dependent plasticity may alter network reorganization. We first show that when plasticity is absent, the Type 1 networks, as expected, exhibit asynchronous dynamics with oscillators of the highest natural frequency robustly evolving faster in terms of their phase dynamics. At the same time, the Type 2 networks synchronize, with oscillators locked so that the ones with higher natural frequency have a constant phase lead as compared to the ones with lower natural frequency. This relationship establishes a robust mapping between the frequency and oscillators’ phases in the network, leading to structure/frequency mapping when plasticity is present. Further we show that while connection plasticity can produce stable synchrony (so called splay states) in Type 1 networks, the structure/frequency reorganization observed in Type 2 networks is not present.

## Introduction

The Kuramoto model of coupled oscillators (1-3) and its variations is often used as a simple way to understand the behavior of coupled neurons in the brain. In this paper, we consider a variation of the model which allows changes in the form of the coupling function to mimic changes in the phase response curve (PRC) (4, 5) of neurons and also to introduce dynamic changes in the coupling motivated by the physiological process of spike-timing dependent synaptic plasticity (STDP) (6, 7). We show that not only does the PRC critically alter network reorganization, but also that plasticity itself affects patterns of network activation. Further, we show that STDP-like plasticity rule together with the modulation of PRC from Type 1 to Type 2 provides dynamic mapping between frequency based information representation where oscillators with higher natural frequency (representing higher external input) evolve faster with Type 1 network coupling, and structural network reorganization based on phase interdependencies established when Type 2 network coupling is present.

The motivation for this particular version of the model is as follows: synchrony of neuronal activity is thought to play crucial role in the brain during healthy function as well as disease (8-10) and is thought to underlie memory formation via network reorganization (7, 11). Also, neuromodulatory effects in the brain may diametrically change network dynamics and resultant network reorganization due to spike-timing dependent synaptic plasticity (STDP). Acetylcholine (ACh) is one of the critical regulators of neural excitability that is essential for brain processes ranging from sleep to cue detection (12-17). Among its various effects, ACh modulates the excitability of neurons by its interaction with the muscarinic receptor system (18). An important downstream target of these signals are slow non-inactivating potassium channels. These channels, and their corresponding ionic current, are blocked when ACh is high and are responsible for a switch of the PRC between integrator and resonator excitability types (19).

ACh modulation of this current exerts continuous control of neuronal excitability properties. On the extremes of this range are two predominant excitability types: Type 1 or Type 2. These two excitability types differ in the dynamical mechanism of spike generation (20-22). One of the characteristics that undergoes the most dramatic change when excitability type is switched is the PRC (4, 5) which measures the differential response to brief and weak stimuli in terms of spike timing perturbation (i.e., advance or delay). Type 1 and Type 2 neurons display significant differences in PRC shape. A Type 1 PRC is uniformly positive, meaning that perturbations will always advance the timing of the next spike. Type 2 neurons have a biphasic PRC, meaning that depending on the timing of the perturbation, it will either advance or delay the next spike. The biphasic character of the Type 2 PRC allows these neurons to synchronize spike firing due to the ability to either shorten or elongate the period, with zero value of phase response becoming a stable fixed point of the dynamics (23).

The differences in propensity for synchronization between high and low ACh states influences network firing dynamics and consequently plasticity patterns regulated by STDP (24, 25). Specifically, robust potentiation (and depotentiation) by STDP relies on consistent relative spike timings between pairs of cells (7). Synchronized network dynamics that are promoted by low ACh modulation support such consistent firing so that most cells will fire within a network burst whose duration is within the STDP time window. Thus, we show that ACh modulation,via changes in PRCs, provides universally effective way of mapping of firing frequency of neurons to structural network representation obtained through STDP - like rule.

## Results

### Equivalence of Type 1/Type 2 PRC Kuramoto model to that with phase delays

We consider a continuous shift from Type 1 to Type 2 PRC on synchrony of networks of non-identical Kuramoto oscillators (1) by introducing a parameter that allows for a continuous shift between experimentally observed (5) Type 1 and Type 2 states (Fig. 1A):

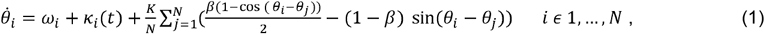

where *β* ∈ [0.1] and is a parameter controlling the transition between the Type 1 and Type 2 (see Fig. 1A). Type 1 dynamics corresponds to *β* = 1 and Type 2 dynamics, *β* = 0 .The noise variable, *k*_*i*_(*t*), is drawn from a Gaussian distribution with μ = 0 and 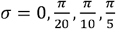. The constant K is the coupling strength, *ω*_*i*_ is the natural frequency of i^th^ oscillator and *θ*_*i*_ denotes its momentary phase. The number of oscillators, *N* = 20. The natural frequencies of the oscillators were distributed according to:

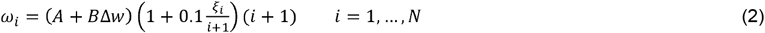

Here A = 0.0125, B = 0.0005, Δ*w* is the natural frequency spread, *ξ*_i_ is a random variable taken from a normal distribution so that the differences in frequency between of i^th^ and j^th^ oscillators fluctuate around (*A* + *B*Δ*w*). This specific shape of the ω_I_ allowed for the frequency spread to be an integer: Δ*w* ∈ [0,9]. Finally, ***i*** is the subscript of the i^th^ oscillator. These natural frequencies loosely represent a cumulative signal coming from sensory modalities external to the modelled network.

**Figure 1.**
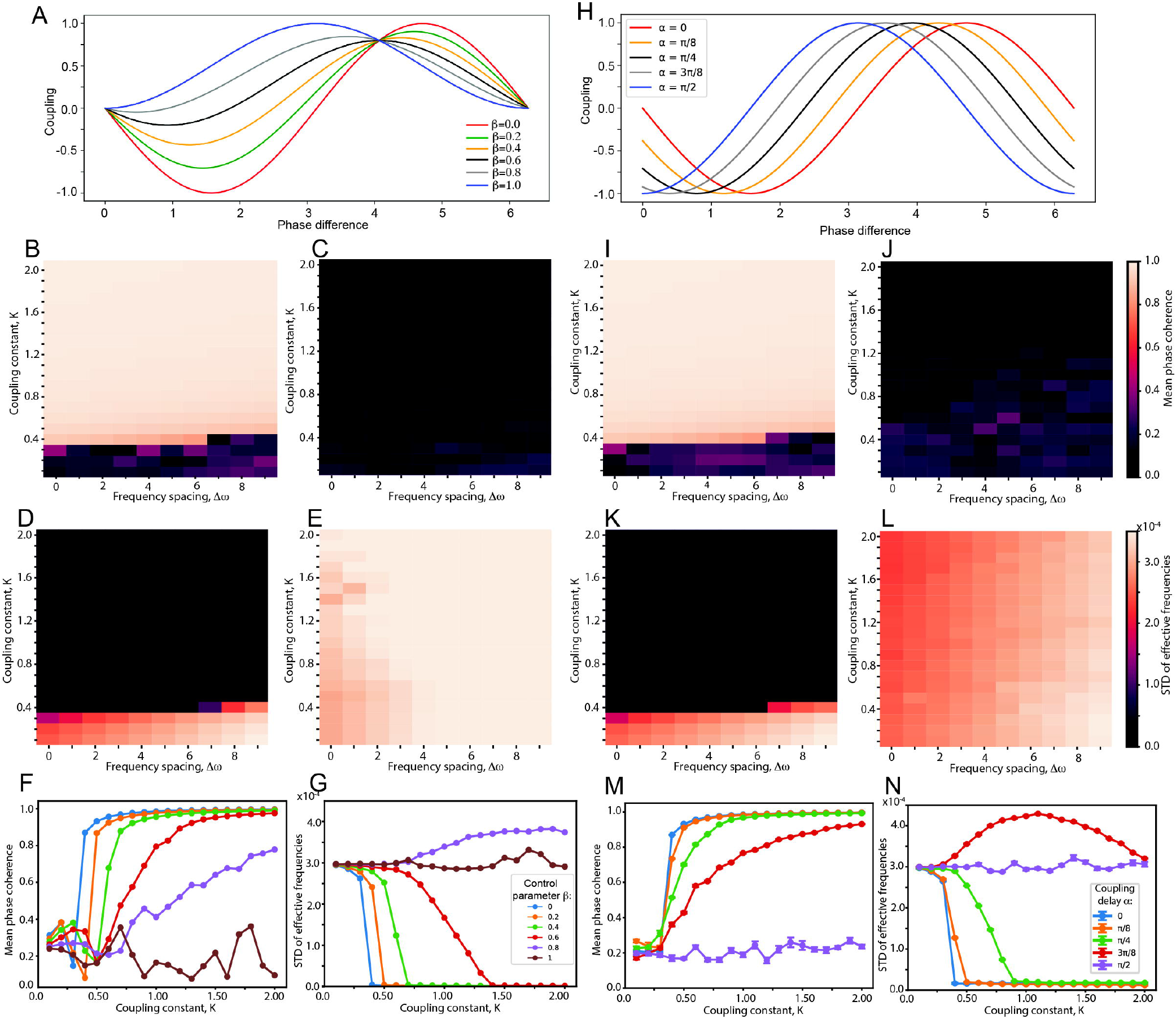
Comparison of dynamics of network of Kuramoto oscillators with different realizations of coupling phase response term. A-G) Phase response curve as described by Eqn. 1: A) PRC as a function of control parameter. *β* B-C) Mean phase coherence as a function of oscillators’ frequency spread (Δ *ω*) and coupling constant K for type 2 (B, *β* = 0) and type 1 (C, *β* = 1) PRC; D-E) standard deviation of observed oscillators frequencies, Ξ, as a function of oscillators’ frequency spread (Δ *ω*) and coupling constant K for type 2 (D, *β* = 0) and type 1 (E, *β* = 1). Mean phase coherence (F) and standard deviation (G) of observed oscillators frequencies, Ξ, as a function of coupling constant for different values of *β*. H-N) same as A-G) but for oscillator coupling having phase delay, *α* (Eqn. 3). In panels F), G), M) and N), the mean difference in natural frequencies between consecutively labeled oscillators ∣ *ω*_*i*+1_ − *ω*_*i*_ ∣ = 0.0145.

We measured the mean phase coherence (26) and standard deviation of observed frequencies, 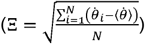 of the oscillators as a function of frequency spread, Δ*w*, and the coupling constant K. We observe two types of states: full synchrony – when the mean phase coherence achieves value of one and standard deviation of observed oscillator frequencies tends to zero and globally asynchronous states - when mean phase coherence tends to zero and the standard deviation of observed oscillator frequencies is non-zero.

As expected, the synchrony changed dramatically between Type 1 and Type 2, with Type 1 networks showing no appreciable synchrony (Fig. 1 B, D, F, G) and Type 2 networks exhibiting a fully synchronous state for wide range of parameters (Fig. 1 C, E, F, G). In the asynchronous state, oscillators with highest natural frequency have the highest observed frequency, while in the synchronized state the oscillators with the higher natural frequency lead (with nonzero phase) those with lower frequency (27).

We compared these results to a network of similar Kuramoto oscillators coupled with phase delays, the Sakaguchi-Kuramoto (SK) model (2) (Fig. 1H – M):

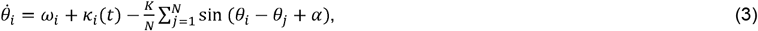

where 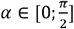 is a phase delay. The results are matching very well (Fig. 1H-M) with Type 2 dynamics corresponding to *α* = 0, and Type 1 dynamics to *α* = *π* /2. Since the SK model has been studied extensively in the literature, we will use it in the remainder of this paper.

The addition of noise does not dramatically change the behavior of the network (Fig. 2) for *α* = *π* /2, as we continue to see asynchronous behavior across different coupling strengths, K.

**Figure 2.**
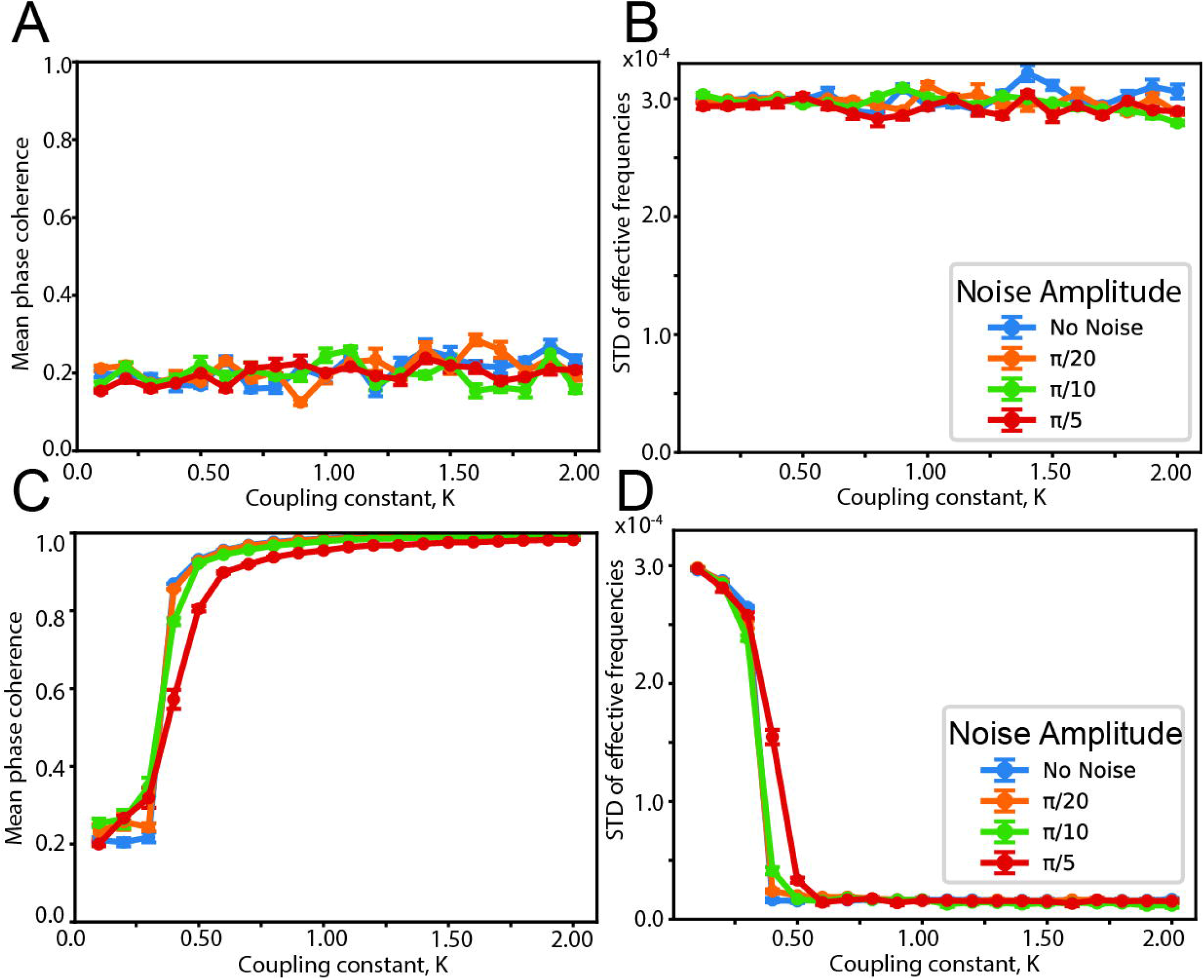
Effects of noise in network consisting of type 1 (*α* = *π* /2) and type 2 (*α* = 0)\oscillator coupling. A, B) Type 1 coupling; Mean phase coherence (A) and standard deviation of observed oscillator frequencies (B) as a function of coupling constant, K, for varying noise levels. C, D) Type 2 coupling – same as A) and B), respectively. The mean difference in natural frequencies between consecutively labeled oscillators ∣ ω_*i*+1_ − *ω*_*i*_ ∣ = 0.0145.

However, for *α* = 0, the transition from asynchronous dynamics to synchronous is increasingly delayed to higher K and not as abrupt as more noise is introduced.

### Effects of spike timing dependent plasticity on network dynamics

Next, we introduce a biologically realistic learning rule mimicking spike timing dependent plasticity rule to the network, with its formulation adopted from (28). The equations now take the form:

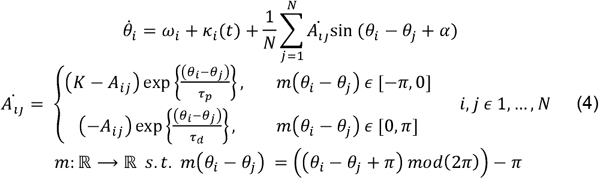

Where *A*_*ij*_ is the coupling strength between i^th^ and j^th^ oscillator, K is the maximal coupling constant that can be achieved in the network; *τ*p = 0.15 and *τ*d = 0.30 denote potentiation and depression time constants, respectively. The plasticity rule mimics a biologically realistic STDP rule between neuronal synapses (7), as it effectively strengthens the connections from leading to following oscillators (in terms of their phase difference), while suppressing the connections from following to leading oscillators, with the major difference that discrete spike timing differences are replaced with continuous measurement of phase differences.

Addition of synaptic plasticity significantly changes the observed global dynamics of the system(Fig. 3). Most significantly, we observe a third type of state emerging for Type 1 networks *α* = (*τ* /2) - the global splay state (29) where phases are locked but randomly distributed (causing mean phase coherence to be near zero) while oscillator frequencies cluster (causing their standard deviation to also tend to zero) (Fig. 3). Generally, this state appears for increasing coupling strength (Fig. 3B, D and Fig. 3F violet line). However, for intermediate values of the coupling constant (K) they can form multiple clusters, while for large K the observed oscillator frequencies converge to the same value (please refer to Fig. 5C, E). The splay state is robust to lower noise values but is largely destroyed with higher noise amplitudes (Fig 4. A, B).

**Figure 3.**
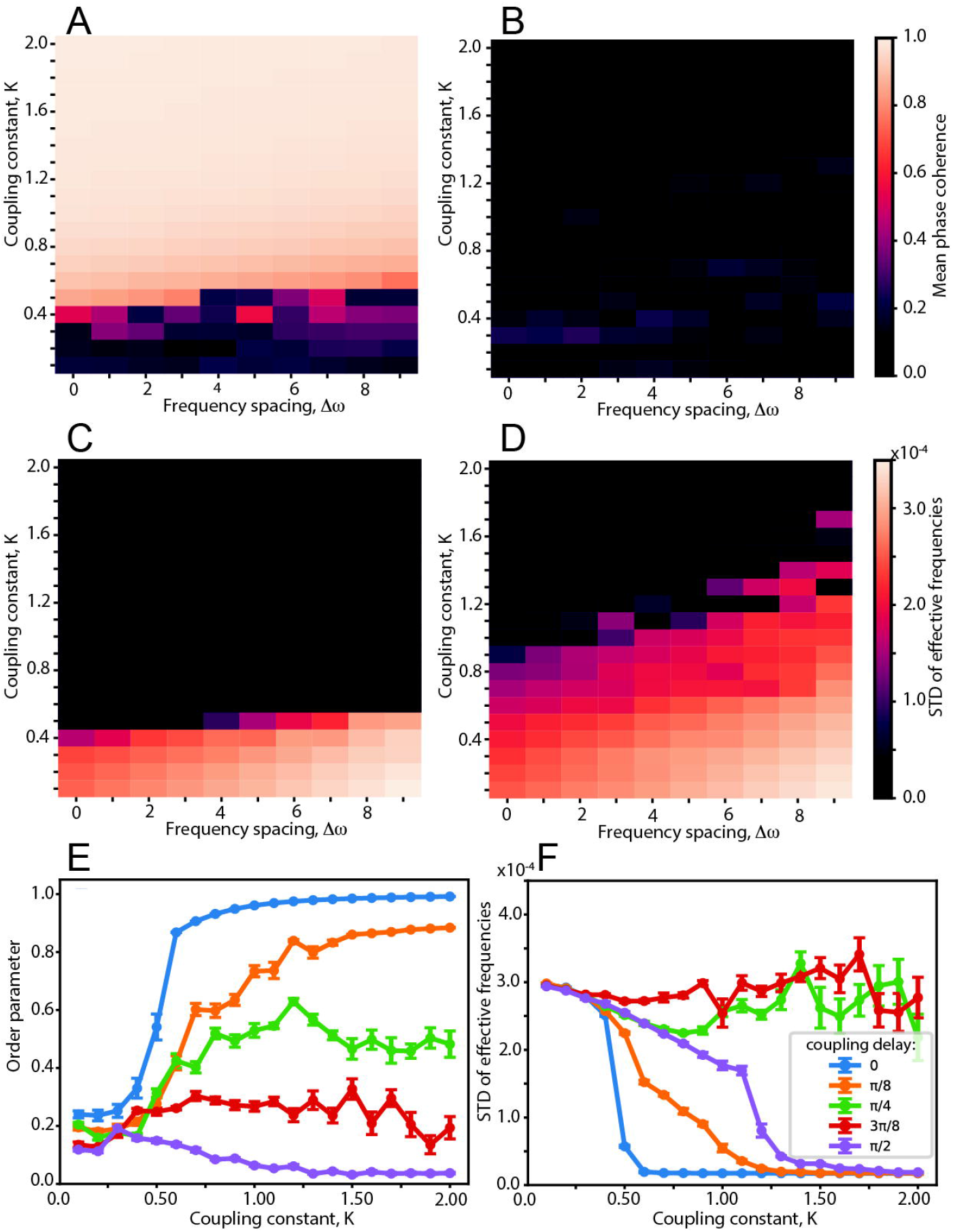
Dynamics of the network with biologically realistic activity dependent connection plasticity. A, C) Type 2 coupling (*α* = 0); Mean phase coherence (A) and standard deviation of observed oscillator frequencies (C) as a function of maximal coupling constant, K. B, D) Type 1 coupling (*α* = *π* /2) ; Mean phase coherence (B) and standard deviation of observed oscillator frequencies (D) as a function of maximal coupling constant, K. E, F) Mean phase coherence (E) and standard deviation of observed oscillator frequencies (F) as a function of maximal coupling constant for different coupling delay *α*. For panels E) and F), the mean difference in natural frequencies between consecutively labeled oscillators ∣ *ω*_*i*+1_ − *ω*_*I*_ ∣ = 0.0145.

**Figure 4.**
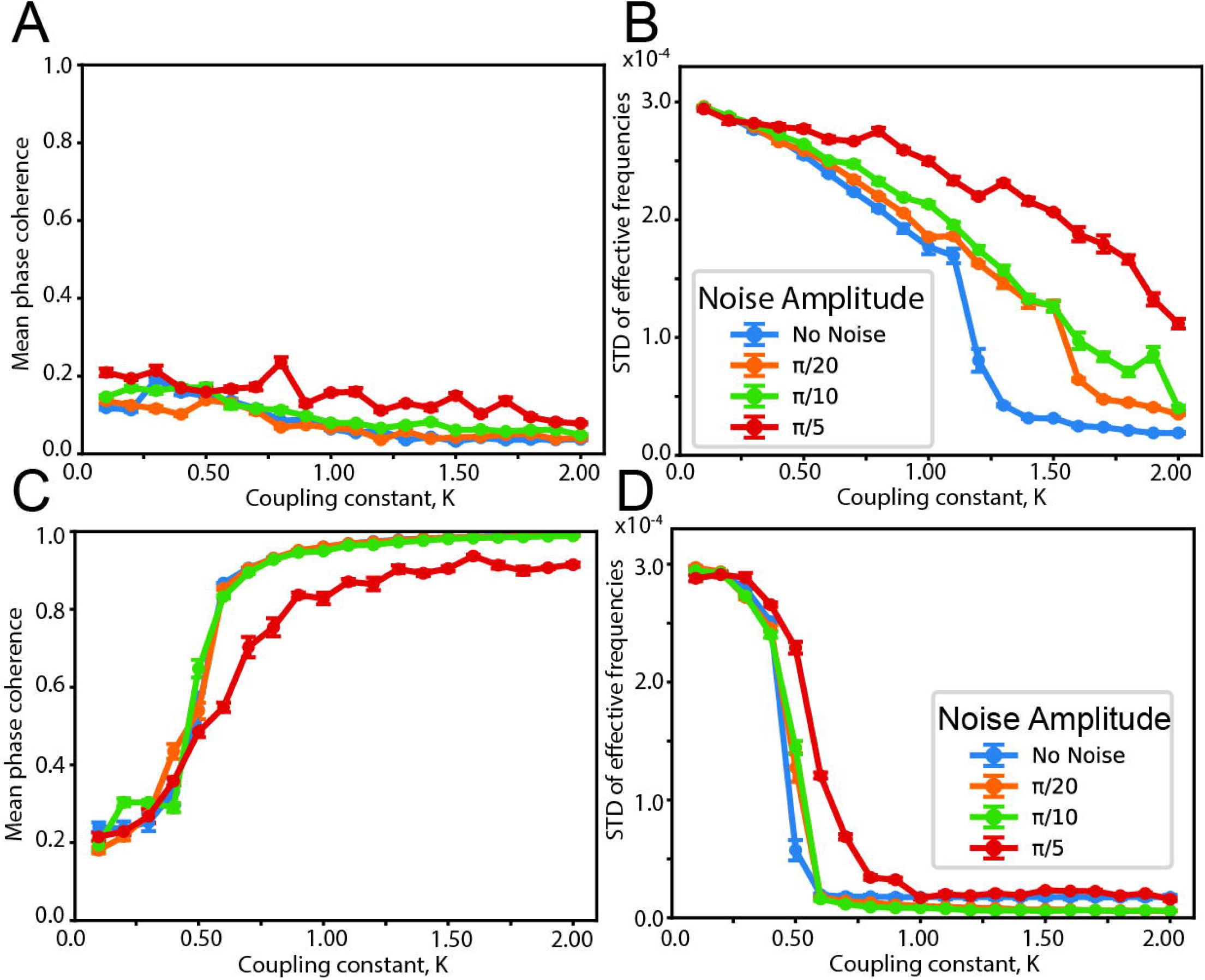
Effects of noise on network dynamics with activity dependent connection plasticity. A, B) Type 1 coupling) (*α* = *π* /2) ; Mean phase coherence (A) and standard deviation of observed oscillator frequencies (B) as a function of maximal coupling, K, for varying noise levels. C, D) Type 2 coupling (*α* = 0); – same as A) and B), respectively. The mean difference in natural frequencies between consecutively labeled oscillators ∣ *ω*_*i*+1_ − *ω*_*i*_ ∣ = 0.0145.

**Figure 5.**
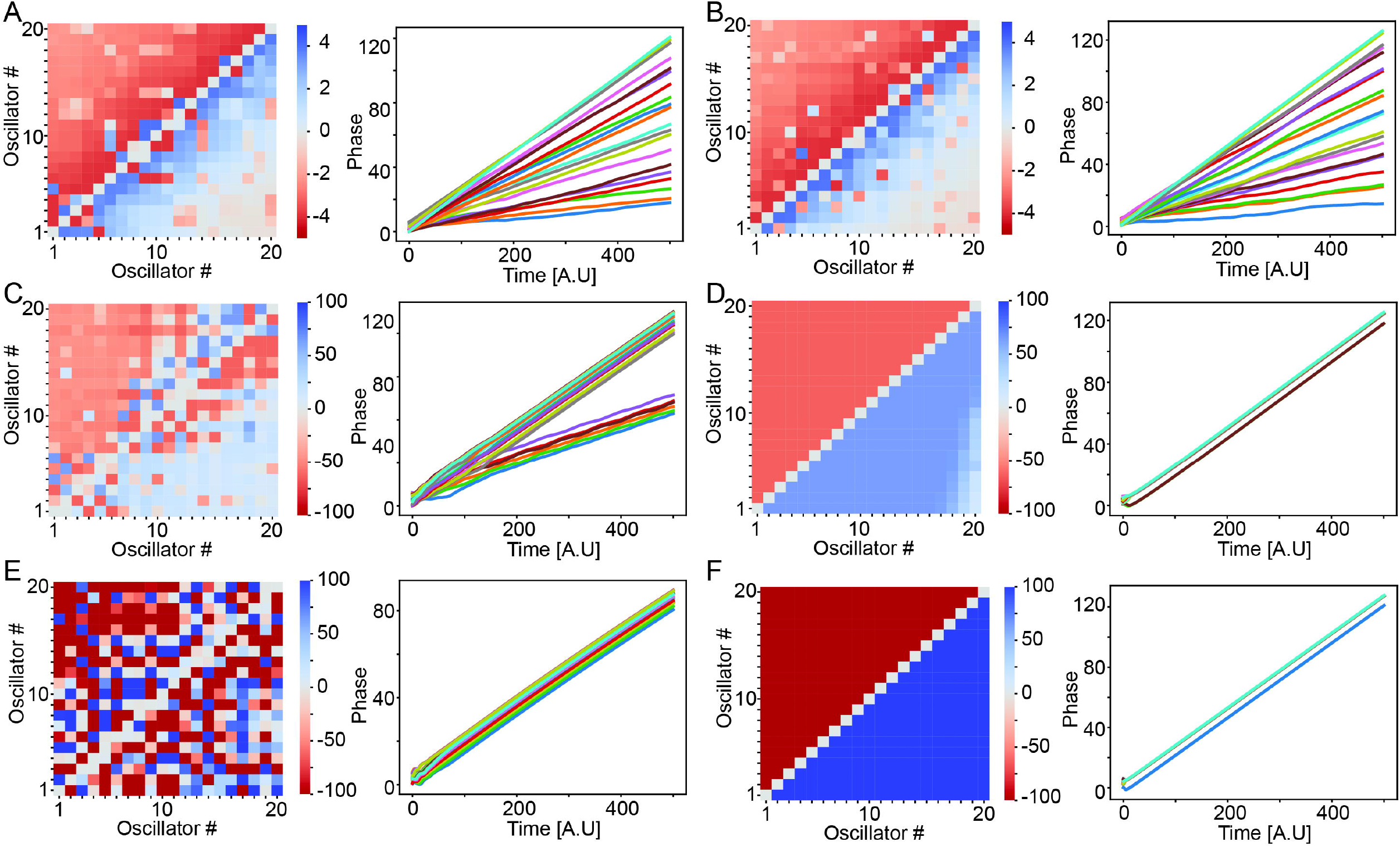
Reorganization of the connectivity matrix and dynamics of the oscillators for type 1 and type 2 network with activity dependent plasticity. A) Type 1 coupling, with maximal coupling K = 0.2 (final reorganization of connectivity matrix – left; evolution of oscillators’ phase as a function of time – right); B) Type 2 coupling with maximal coupling K = 0.2; C) Type 1 coupling with maximal coupling K = 0.8; D) Type 2 coupling with maximal coupling K = 0.8; E) Type 1 coupling with maximal coupling K = 1.6; F) Type 2 coupling with maximal coupling K = 1.6. For all runs, the mean difference in natural frequencies between consecutively labeled oscillators ∣ *ω*_*i*+1_ − *ω*_*i*_ ∣ = 0.0125.

Addition of plasticity to Type 2 dynamics, on the other hand, causes the transition from asynchronous-to-synchronous dynamics to happen at higher values of the coupling constant K, but, qualitatively, this transition does not change (Fig. 3A, C, E). The reason for this delayed transition (in terms of K value) is the fact that the plasticity rule effectively disconnects about half of the synapses in the network - reducing overall connectivity (see Figs. 5 and 6 for reference). With plasticity present, addition of noise (Fig. 4C, D) also has a bigger effect on the quality of synchronization (as compared to the case when plasticity is absent) for the same reason.

**Figure 6.**
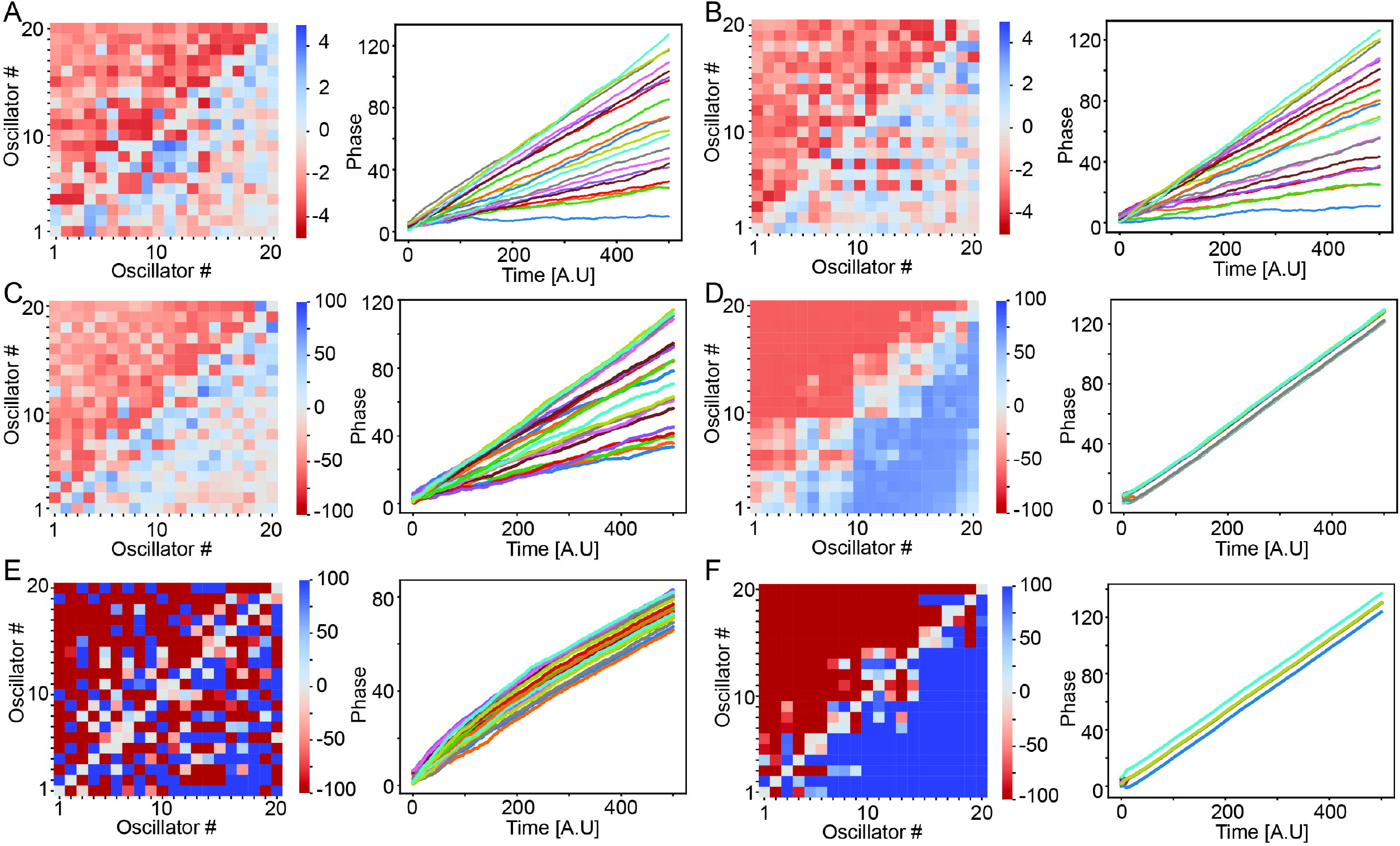
Reorganization of the connectivity matrix and dynamics of the oscillators for type 1 and type 2 network with activity dependent plasticity, in presence of noise. A) Type 1 coupling, with maximal coupling K = 0.2 (final reorganization of connectivity matrix – left; evolution of oscillators’ phase as a function of time – right); B) Type 2 coupling with maximal coupling K = 0.2; C) Type 1 coupling with maximal coupling K = 0.8; D) Type 2 coupling with maximal coupling K = 0.8; E) Type 1 coupling with maximal coupling K = 1.6; F) Type 2 coupling with maximal coupling K = 1.6. For all runs, the mean difference in natural frequencies between consecutively labeled oscillators ∣ *ω*_*i*+1_ − *ω*_*i*_ ∣ = 0.0125. and noise variance *σ* = *π*/10,

### Differential structural network reorganization

Finally, we investigated network structural reorganization for both the Type 1 and Type 2 PRC as a function of maximal coupling constant (K). Figure 5 depicts these results for the no noise case. The color maps depict changes in the adjacency matrix of the networks. The oscillators are ordered with respect to their natural frequencies, with the slowest oscillator positioned in bottom left corner. The network is initially coupled with *A*_*ij*_ = *K*/2, thus change in connection of negative 100% means complete disconnection of the two oscillators, while a change of 100% means convergence of the given connection to its maximal possible value (K). The graphs positioned to their right show examples of evolution of their phase angles as a function of time. For low values of K (i.e., K=0.2), we observe minimal network reorganization (note color scale) for both Type 2 (*α* = 0) and Type 1 (*α* = *π*/2) PRC (Fig. 5A, B), with oscillators lying next to diagonal exhibiting the strongest reorganization. This is due to the fact that only oscillators with very similar frequencies can exhibit persistently synchronized activity, leading to strengthening or weakening of the coupling. The difference becomes more pronounced for intermediate and high values of the maximal coupling. When K=0.8, we observed the formation of splay states for the Type 1 PRC, while the network with Type 2 oscillators displayed full synchrony with the faster oscillators leading the slower ones as expected (Fig. 5C, D) (27).

Splay states are characterized by locking of the phases at large phase differences, leading to zero phase coherence. The connectivity pattern still follows the general trend of faster oscillators leading the slower ones but with additional clusters forming with reverse dependence (Fig. 5C). The example on the right of Fig. 5C shows the formation of two splay clusters with two different mean observed frequencies of the oscillators. The evolution of the two clusters will lead to their structural disconnection and fragmentation of the network.

Finally, for the strongest maximal coupling (K=1.6) we observe a global splay state forming for the network consisting of Type 1 oscillators with no clear structure/natural frequency dependence. However, the network consisting of Type 2 oscillators is fully synchronized with pronounced frequency-structure dependence.

When noise is present (Fig. 6), the weakest coupling exhibits minimal changes in the connectivity matrix (note the scale) for both networks consisting of Type 1 and Type 2 oscillators (Fig. 6A, B). The splay state for the intermediate value of maximal coupling (K=0.8) is largely destroyed (Fig. 6C). The network consisting of Type 2 oscillators exhibits global organization (Fig. 6D), with oscillators of the most similar frequencies (those lying near the diagonal) exhibiting ordering shifts. This pattern occurs because these oscillators have the smallest phase differences when they synchronize – thus, noise can easily alter this ordering. We observe minimal changes in ordering due to noise for the highest maximal coupling (K=1.6): the network consisting of Type 1 oscillators achieves a global splay state with no structure/phase ordering (Fig, 6E). The network consisting of Type 2 oscillators is largely synchronized with strong frequency-coupling ordering, except for few small clusters forming near diagonal.

## Discussion

We have shown that networks consisting of non-identical Type 1 or Type 2 Kuramoto oscillators can lead to significantly different dynamics and patterns of structural network reorganization when a biologically realistic plasticity rule is applied. Type 2 networks exhibit global synchrony for a large parameter space, with global frequency-structure dependence so that connections from faster oscillators to slower oscillators are strengthened and the reverse connections are weakened. Type 1 networks, as predicted, do not exhibit robust synchrony for most parameter values, with a new type of state – the splay state - emerging when the network exhibits reorganization with plasticity (29). Here, the oscillators have locked frequencies, but their phase differences remain random. Those splay states are more vulnerable to noise and do not exhibit a clear frequency-structure relationship as the Type 2 global synchrony does.

These results are of considerable relevance to brain function since neuromodulatory effects of M1 ACh receptors change the neuronal PRC from Type 2 (low ACh, for example NREM sleep) to Type 1 (high ACh; waking behavior and REM sleep) (4, 5, 12, 30). In this case, the natural frequency of the oscillators would represent the input from external brain modalities to the network. When the plasticity is absent, this information is preserved in the networks via relative observed frequency in Type 1 networks and phase delay between the Type 2 oscillators in the synchronized state. Switching between Type 1 and Type 2 (via ACh modulation) provides consistent mapping between phase relationships (in Type 2 network) and frequency relationships (in Type 1 network).

The situation is significantly different when coupling dynamics (via STDP) are introduced, as the splay state emerges for a strongly coupled Type 1 network. The global synchrony for Type 2 networks exhibit global structural mapping of the natural frequencies of the oscillators onto connectivity space, with faster oscillators strongly coupling to slower oscillators, while the reverse connection is silenced. For the Type 1 oscillators in the splay state, this mapping is not present.

This suggests that Type 2 networks can create a structural snapshot of the relative intrinsic properties of the oscillators, while the Type 1 network cannot, leading to information loss. We hypothesize that the differences in observed dynamics, and the formation of splay states may represent the difference between brain activity during epileptic episodes and normal cognitive function. Highly synchronous seizure dynamics would correspond to a splay state where the structure/phase relationship is abolished leading to temporary cognitive deficits.

## Acknowledgements

This work was supported by grant NIH MH135565 (MZ).

